# Lipid constituents from *Cissus trifoliata* stems

**DOI:** 10.1101/2023.09.27.559825

**Authors:** Luis Fernando Méndez-López, José Luis González Llerena, Bryan Alejandro Espinosa-Rodríguez, Isaías Balderas-Rentería, María Del Rayo Camacho-Corona

## Abstract

**Objective:** The stems from plants of the genus *Cissus* are used in traditional medicine worldwide. In Mexico *Cissus trifoliata* is employed for the management of several diseases, however little is known about the active compounds. The present investigation focuses on the extraction, isolation, and characterization of the lipid contents from the stems of *C. trifoliata* by chromatographic, spectroscopic, and spectrometric techniques.

**Methods:** The hexane extract was fractionated sequentially with hexane and ethanol and visualized by TLC. The compounds were isolated by column chromatography and structurally elucidated using NMR. In addition, the complete extract and solids obtained by fractionations were analyzed by GC-MS.

**Results:** According to NMR signals, solids present in the hexane extract correspond to hentriacontane, octatetracontane, octacosanoic acid, tricosyl tetracosanoate and sitosterol. Furthermore, the analysis by GC-MS found 33 compounds, and the most abundant were hentriacontane, squalene, palmitic acid, oleic acid, β-sitosterol, and nonacosane.

**Conclusion:** This report describes the major constituents in the hexane extract from *Cissus trifoliata* stems. Our analysis suggests an abundant composition of higher alkanes, fatty acids, and sterols, which are the major lipid components of cuticular waxes and plant cell membranes.

## Introduction

*Cissus trifoliata* (L.) ^1^ also known as “*Hierba del buey*” is widely distributed in the Mexican territory ^2^ and documented as an important medicinal plant for the management of several diseases ^3^. Ethnobotanical uses for *C. trifoliata* in Mexico include gastrointestinal illnesses ^3^, sores ^4^, skin infections, inflammation, abscesses, and tumors ^5^. Recent analysis indicates a metabolic profile of *C. trifoliata* stems enriched in terpenes, flavonoids, and stilbenes ^6^. Namely, forty-six metabolites were identified and include alcohols, alkanes, esters, fatty acids, terpenes, and phenolic compounds ^6^. In addition, the analyzed extracts showed cytotoxic activity against cancer cell lines from the liver, breast, prostate, cervix, and lung ^6^. Furthermore, the bioassay-guided study of the chloroform-methanol extract determined a fraction that reduces the 80% of proliferation on prostate cancer cells (PC3) ^7^. According to HPLC-QTOF-MS analysis, the antiproliferative effects could be attributed to the presence of coumaric and isoferulic acids, apigenin, ursolic and betulinic acids, hexadecadienoic and octadecadienoic fatty acids and the stilbene resveratrol ^7^. However, few reports describe the lipid constituents from the stems of *Cissus* plants. Moreover, the isolation of compounds from *Cissus* plants had centered in the aerial parts or the leaves ^8-10^ using polar solvents such as acetate, ethanol, methanol, or hydroalcoholic extractions ^11, 12^. At present, only one report of the isolation and structural elucidation of compounds in the hexane extract of the stems of *Cissus quadrangularis* has been published. They found 29 compounds, that belong to the chemical classes of triterpenes, fatty acids, methyl esters, glycerolipids, steroids, phytols, and cerebrosides. The most common type of isolated compounds were the octadecenoic acid methyl esters, the 3-O-linoleoyl glycerol and derivatives, beta-sitosterol, dammara-type and lupane triterpenes, and phytols ^13^. In another report that includes the lipid contents of the stems (hexane extract of aerial parts) of *C. quadrangularis* were isolated the alcohols, esters, fatty acids, sterols, long chain alkanes, ketones, sterols, and triterpenes. Specifically, the 4-hydroxy-2-methyl-tricos-2-en-22-one, 9-methyl-octadec-9-ene, heptadecyl octadecanoate, icosanyl icosanoate, 31-methyltritriacontan-l-ol, 7-hydroxy-20-oxo-docosanyl cyclohexane, 31-methyltritriacontanoic acid, friedelan-3-one, taraxerol and isopentacosanoic acid and b-sitosterol ^14^. In this contribution we report the extraction, chromatography, NMR, and GC-MS analysis of lipid constituents in the hexane extract from the stems of *C. trifoliata*.

## Materials and methods

### General

Column chromatography (CC) was carried out on silica gel (EMD chemicals Inc.). Thin layer chromatography (TLC) was performed on 60 F254 silica gel 0.2 mm on aluminum plates (Merck). TLC plates were observed under UV lamp (Spectroline λ 254 nm and 365 nm) and revealed with ceric sulfate solution. Determination of melting point (MP) was performed on a Fischer-Johns apparatus. One dimensional Nuclear Magnetic Resonance (NMR) spectra were obtained using a Bruker AVANCE III HD 400 MHz instrument, with deuterated solvents, and tetramethylsilane (TMS) as internal standard. The software employed for the processing of NMR spectral data was Mestrenova version 12. Finally, GC-MS analysis of hexane extract was performed on a GC-MS Agilent 6890, MSD 5973N using the database of the National Institute Standard and Technology (NIST) having more than 62,000 patterns version 1.7A.

### Plant material and extraction

*Cissus trifoliata* was collected in Rayones Nuevo León, Mexico, and identified by a trained biologist. A reference sample (Voucher number 027499) was deposited in the Department of Botany, of the Universidad Autonoma de Nuevo Leon. Dried and ground stems (756 g) were extracted by maceration with hexane (1.5 L), yielding 3.5 g of dried extract. The organic extract was filtered and concentrated under reduced pressure using a rotary evaporator at 40 °C. The extract was kept at 4 °C until used.

### Isolation and characterization of compounds from the hexane extract

The hexane extract was subjected to column chromatography, using silica gel (70 g) as the stationary phase and hexane-ethanol in gradient as the mobile phase. A total of 224 fractions of 20 mL each were collected and pooled according to their chromatographic similarity in four subfractions (I-IV). A white amorphous solid precipitated (50 mg) from fraction I (1-84, hexane 100%). The precipitated solid was separated by filtration and washed with cold acetone to remove soluble impurities. TLC analysis of the resultant white, waxy solid showed no activity under visible or UV light but stained with cerium sulfate as a brown spot. From fraction II (F85-98, hexane-ethanol 96:4) a cream amorphous solid precipitated. The solid was soluble in hexane and insoluble in acetone, thus, it was filtrated and washed with cold acetone. A white solid was recovered (28 mg) and was visualized as a brown spot in a TLC stained with cerium sulfate. The rest of the fraction II was passed through a column chromatography of silica gel (2.35g) using a gradient of hexane-ethyl acetate. A white waxy solid (39 mg) precipitated from subfractions F30-37 (hexane-ethyl acetate 90:10) and washed with methanol to remove impurities. The compound was soluble in hexane and acetone, showing a brown spot in TLC stain with cerium sulfate. From the subfractions F56-63 (hexane-ethyl acetate 85:15) precipitated a solid that was purified by successive recrystallization with acetone giving 35 mg of needle shape crystals, that were soluble in chloroform and acetone. The TLC staining with cerium sulfate resulted in a purple spot. From the *mother* liquor of which compound 3 was obtained a white solid (30 mg) was further vacuum-filtered and washed with hexane. The chloroform soluble compound was visualized as a brown spot in a TLC stained with cerium sulfate. The pooled fractions III (F99-144, hexane-ethanol 93:7), and IV (F145-224, ethanol 100%) were chromatographed but no solids were obtained.

### Gas chromatography-mass spectrometry analysis

The hexane extract and the isolated compounds were examined by GC-MS to determine their chemical composition. The analysis was conducted with the column HP-5 (30 mm × 0.25 mm × 0.25 μm). The carrier gas was helium with a gas flow rate of 1mL/min and a linear velocity of 37cm/s. The injector temperature was set at 270°C. The initial oven temperature was 70°C and programmed to increase to 310°C. The program was 70° to 200°C in 10°C/min, 200°C to 310°C at 10°C/min and the final temperature was held for 5 min at 310°C. The mass spectrometer was operated in the electron ionization mode at 70 ev and electron multiplier voltage at 1859 V. The compounds were identified by comparison of the spectral data and fragmentation pattern.

## Results and Discussions

### Characterization of compounds from the hexane extract of *C. trifoliata* stems

Compound 1 was a white, waxy solid, with a melting point of 60-62°C. The ^1^H NMR (400 MHz, CDCl_3_) spectra of the compound showed a triplet at δ 0.88 (t, J = 6.8 Hz, H1, H31) integrating for six methyl protons and a singlet of methylenes at δ 1.25 (H2 to H30) integrating for 58 protons which suggest a long chain linear alkane. The ^13^C NMR spectra show five carbon signals at δ 31.95 (C3, C29), 29.72 (C5 to C27), 29.38 (C4, C28), 22.71 (C2, C30), 14.12 (C1, C31), which corresponds to the observed five resonances for a hydrocarbon chain of (αCH3)_2_(βCH2)_2_(γ CH2)_2_(δ CH2)_2_(ε CH2)_2._ The terminal αCH_3_ groups, and four CH_2_ groups (β, γ, δ, and ε), at 14.3 ppm (αCH3), 23.0 ppm (βCH2), 32.2 ppm (γCH2), 29.7 ppm (δ CH2) and 30.0 ppm (ε CH2) ^15^. Thus, compound 1 was characterized as hentriacontane, which is a major constituent of plant cuticular waxes ^16^, but is the first time isolated in a plant of the genus *Cissus*.

Compound 2 was recovered as a white solid with a melting point of 74-75°C. The ^1^H NMR (400 MHz, CDCl_3_) spectra showed a triplet at δ 0.88 (t, J = 6.6 Hz, H1, H48) integrating for six methyl protons and a singlet of methylenes at δ 1.25 (H2 to H47) integrating for 92 protons. The ^13^C NMR spectra shows five carbon signals at δ 31.95 (C3, C46), 29.72 (C5 to C43), 29.38 (C4, C45), 22.71 (C2, C47), 14.12 (C1, C48). Thus, compound 2 corresponds with octatetracontane, which is a common constituent in plant stems ^17-19^, but is the first time isolated from the genus *Cissus*.

Compound 3 is a white waxy solid with a melting point of 64-65°C. The ^1^H NMR spectral data shows a triplet at δ 0.88 (t, J = 6.7 H, H28) attributed to three methyl protons, and at 1.25 (H4 to H27) a singlet integrating for 48 protons, which corresponds to 24 units of methylenes, downfield at 1.63 (H3) a multiplet integrating two protons corresponding to a methylene protons beta, in respect to a carbonyl, and at 2.35 (t, J = 7.5 Hz, H2) a triplet integrating for two protons corresponding to an alpha carbonyl methylene protons. The ^13^C NMR data suggests the structure of a carboxylic acid with the molecular formula of C_28_H_56_O_2_, which corresponds with the octacosanoic acid. The signals were assigned; accordingly, at δ 178.47 a signal that corresponds to a carboxylic carbon at C1, along with the signals at 33.79 (C2) of the alpha carbonyl methylene, and 31.94 (C26), 29.69 (C4 to C25), 24.72 (C23), 22.70 (C27) and 14.12 (C1). This is the first report of the presence of octacosanoic acid in a plant from the genus of *Cissus*, although is one of the most common plant wax fatty acids ^20, 21^.

Compound 4 is a needle-shaped crystal with a melting point of 135-136°C. NMR spectral data suggest a mixture of phytosterols. The ^1^H NMR shows the characteristic signals of β-sitosterol plus less intense signals at δ 5.15 (dd, J = 9, 15 Hz), and 5.01 (dd, J = 9, 15 Hz) suggesting the presence of two additional olefinic protons corresponding to stigmasterol. The main signals correspond to β-Sitosterol: six methyl signals that appeared at δ 1.01 (s, 3H, H19), 0.92 (d, J= 6.5 Hz, 3H, H21), 0.86 (t, J = 7.3 Hz, 3H, H29), 0.84 (d, J = 7.2 Hz, 3H, H26), 0.81 (d, J = 7.2 Hz, 3H, H27), and 0.68 (s, 3H, H18). Moreover, the spectra also showed two more characteristic signals of the sterol rings, the olefinic proton in ring B at δ 5.53 (psd, J = 4.7Hz, 1H, H6) and the proton in ring A adjacent to the hydroxyl group at 3.53 (m, 1H, H3). The two additional signals at δ 5.15, and 5.01 corresponds to the double bond between C22 and C23 in the side chain of stigmasterol. The ^13^C NMR showed twenty-nine carbon signals including the characteristic six methyls, the two olefinic carbons, and the carbon adjacent to the hydroxyl group of β-Sitosterol. The assigned carbons were, at δ 140.78 (C5), 121.73 (C6), and 71.83 (C3) for the ring motifs. The methyls were assigned as: at δ 21.10 (C26), 19.83 (C19), 19.41 (C27), 19.05 (C21), 12.00 (C29), and 11.87 (C18). Thus, the spectroscopic data of compound 4 suggest the presence of two sterols, according to the reported spectra in the literature for β-sitosterol and stigmasterol ^22^. β-sitosterol and stigmasterol were previously isolated from the hexane extract of stems of *C*. *quadrangularis* ^13^ and also from methanolic and ethyl acetate extracts of aerial parts and roots of *C. assamica* ^23^, *C. polyantha* ^24^, *C. rheoifolia* and *C. pteroclada* ^25, 26^. β-sitosterol is the principal sterol in plant materials, in addition to its 22-dehydro analogue stigmasterol, they occur in most plants playing roles in the regulation of membrane fluidity, and cellular developmental processes acting as precursors of the brassinosteroids; a class of plant hormones ^27^.

Compound 5 precipitated from the *mother* liquor of compound 3, as a white solid with a melting point of 62-63 °C. The ^1^H NMR shows the presence of two triplets integrating for two protons each, at δ 4.05 (t, J= 6.7 Hz, H25), and 2.29 (t, J = 7.5 Hz, H2), suggesting the presence of methylene groups adjacent to an ester group, and a multiplet at δ 1.60 (H3, H26) integrating for 4 protons corresponding for a couple of methylenes beta position relative to the ester, a singlet at 1.25 (H4 to 23, and H27 to H45) integrating for 80 protons corresponding to a long methylene chain, and a triplet at δ 0.88 (t, J = 6.7 Hz, 6H) for six methyl protons (H24, H47), thus indicating the presence of a long chain saturated ester with a molecular formula of C_47_H_94_O_2_. The ^13^C NMR shows the characteristic carbonyl at δ 174.04 (C1), and the carbon alpha in respect to the oxygen of the ester at δ 64.41 (C25), subsequently appear the signal of the carbon alpha to the carbonyl at δ 34.44 (C2), and a signal at 31.94 (C22, C45) corresponding to the gamma carbons respect to the terminal methyls, then, appears the carbon chain signal at 29.71 (C27 to C44, and C27 to C21). After the methylene chain appears at 25.95 (C26), 25.05 (C3), 22.70 (C46, C23), and 14.12 (C24, C47). Thus, the spectroscopic data suggest the compound tricosyl tetracosanoate; one of the major constituents of the long esters in plant waxes ^28^, although, at present no previous report in a plant of the genus *Cissus* was done.

**Figure 1.**
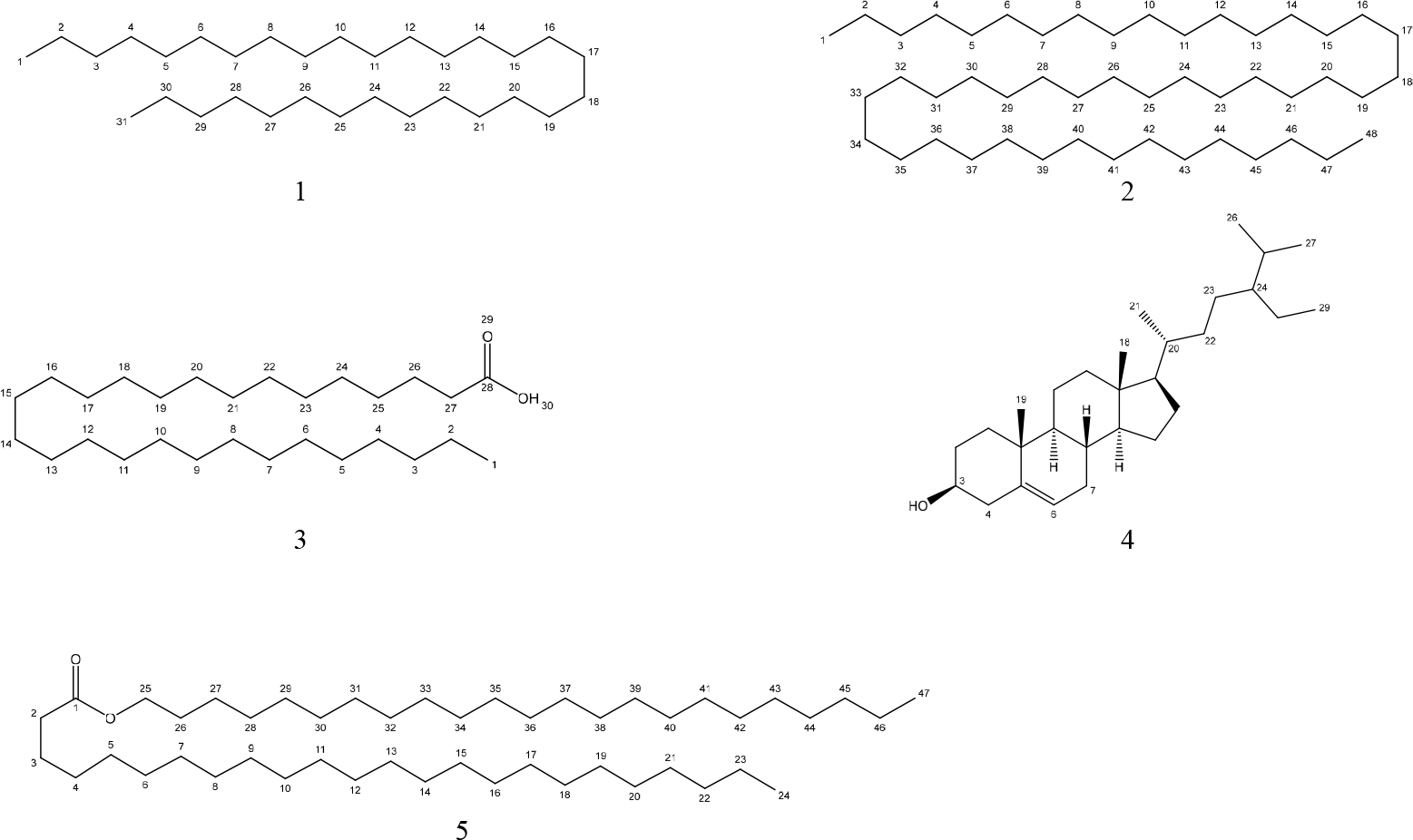
Chemical structures of compounds present in the hexane extract of *C. trifoliata* stems according to spectroscopic analysis

### Identification of compounds analyzed by GC-MS

The chemistry of the lipid contents in the hexane extract of *C. trifoliata* stems was analyzed by GC-MS. The constituents were identified by comparison with the NIST database version 1.7A. The name, retention time, molecular weight, and molecular formula of the components of the extract are listed in table 1.

**Table 1.** Metabolites identified by GC-MS in the hexane extract of *C. trifoliata* stems.

|  | Compound | Area | RT | Abundance (%) | Mass | Formula |
| --- | --- | --- | --- | --- | --- | --- |
| 1 | n-hexadecanoic acid | 1821015375 | 66.472 | 8.222 | 256.430 | C<sub>16</sub>H<sub>32</sub>O<sub>2</sub> |
| 2 | n-hexadecanoic acid ethyl ester | 673724131 | 66.623 | 3.042 | 284.484 | C<sub>18</sub>H<sub>36</sub>O<sub>2</sub> |
| 3 | heneicosane | 218951311 | 71.425 | 0.989 | 296.583 | C<sub>21</sub>H<sub>44</sub> |
| 4 | (z z)-9,12-octadecadienoic acid | 1604487843 | 74.039 | 7.244 | 280.450 | C<sub>18</sub>H<sub>32</sub>O<sub>2</sub> |
| 5 | (z)-9-Octadecenoic acid | 586274008 | 74.328 | 2.647 | 284.480 | C<sub>18</sub>H<sub>34</sub>O<sub>2</sub> |
| 6 | octadecanoic acid | 552786315 | 75.294 | 2.496 | 284.480 | C<sub>18</sub>H<sub>36</sub>O<sub>2</sub> |
| 7 | docosane | 61476191 | 75.754 | 0.278 | 310.610 | C<sub>22</sub>H<sub>46</sub> |
| 8 | tricosane | 557611582 | 80.129 | 2.518 | 324.637 | C<sub>23</sub>H<sub>48</sub> |
| 9 | eicosanoic acid | 253505434 | 83.176 | 1.145 | 312.538 | C<sub>20</sub>H<sub>40</sub>O<sub>2</sub> |
| 10 | pentacosane | 38477994 | 87.794 | 0.174 | 352.691 | C<sub>25</sub>H<sub>52</sub> |
| 11 | octacosane | 913386337 | 88.149 | 5.204 | 394.772 | C<sub>28</sub>H<sub>58</sub> |
| 12 | bis(2-ethylhexyl) phthalate | 399666030 | 89.601 | 1.805 | 390.564 | C<sub>24</sub>H<sub>38</sub>O<sub>4</sub> |
| 13 | 1-hexacosene | 54683947 | 91.447 | 0.247 | 364.702 | C<sub>26</sub>H<sub>52</sub> |
| 14 | pentatriacontane | 17988546 | 91.532 | 0.081 | 492.961 | C<sub>35</sub>H<sub>72</sub> |
| 15 | heptacosane | 195624019 | 95.269 | 0.883 | 380.745 | C<sub>27</sub>H<sub>56</sub> |
| 16 | bis (2-ethylhexyl) isophthalate | 245440442 | 96.721 | 1.108 | 390.564 | C<sub>24</sub>H<sub>38</sub>O<sub>4</sub> |
| 17 | cyclooctacosane | 31056794 | 98.481 | 0.331 | 392.756 | C<sub>28</sub>H<sub>56</sub> |
| 18 | oxyrane heptadecil | 51278173 | 99.854 | 0.232 | 282.512 | C<sub>19</sub>H<sub>38</sub>O |
| 19 | squalene | 2658224840 | 99.941 | 12.002 | 410.73 | C<sub>30</sub>H<sub>50</sub> |
| 20 | nonacosane | 1322290997 | 102.377 | 5.970 | 408.799 | C<sub>29</sub>H<sub>60</sub> |
| 21 | epicholesterol | 17018543 | 106.495 | 0.077 | 386.654 | C<sub>27</sub>H<sub>46</sub>O |
| 22 | brassicasterol | 20058125 | 107.986 | 0.091 | 398.675 | C<sub>28</sub>H<sub>46</sub>O |
| 23 | nonadecane,9-methyl | 1993626 | 108.426 | 0.009 | 282.329 | C<sub>20</sub>H<sub>42</sub> |
| 24 | dotriacontane | 30282752 | 108.433 | 0.137 | 450.88 | C<sub>32</sub>H<sub>66</sub> |
| 25 | hentriacontane | 4966534002 | 109.241 | 22.424 | 436.853 | C<sub>31</sub>H<sub>64</sub> |
| 26 | campesterol | 128043977 | 111.546 | 0.578 | 400.691 | C<sub>28</sub>H<sub>48</sub>O |
| 27 | stigmasterol | 241246607 | 112.571 | 1.089 | 412.702 | C<sub>29</sub>H<sub>48</sub>O |
| 28 | 1,30-triacontanediol | 220025049 | 113.143 | 0.993 | 452.464 | C<sub>30</sub>H<sub>62</sub>O<sub>2</sub> |
| 29 | β-sitosterol | 1421840477 | 114.588 | 6.420 | 414.718 | C<sub>29</sub>H<sub>50</sub>O |
| 30 | tritriacontane | 1019622149 | 114.804 | 4.604 | 464.907 | C<sub>33</sub>H<sub>68</sub> |
| 31 | lupeol | 827035533 | 116.401 | 3.734 | 426.729 | C<sub>30</sub>H<sub>50</sub>O |
| 32 | 1-triacontanol | 77657878 | 117.885 | 0.351 | 438.825 | C<sub>30</sub>H<sub>62</sub>O |
| 33 | stigmast-4-en-3-one | 637225122 | 118.246 | 2.877 | 412.702 | C<sub>29</sub>H<sub>48</sub>O |

The most abundant class of metabolite identified by GC-MS in hexane extract of *C. trifoliata* stems were alkanes (40%), fatty acids (23%), terpenes (20%), fatty alcohols (11%), esters (3%) and phthalates (3%). The major compounds were the hentriacontane, squalene, palmitic acid, oleic acid, β-sitosterol, and nonacosane. Overall, the constituents identified in the hexane extract were consistent with the composition of cuticles and membranes of plants, in which lipid constituents are C16 and C18 esterified fatty acids and waxes (mixtures of homologous series of long-chain aliphatics, such as alkanes, alcohols, aldehydes, fatty acids and esters, together with variable amounts of triterpenes) glycerolipids, sterols, and sphingolipids ^29^. Previous reports of the GC-MS analysis of the hexane extracts of *C. quadrangularis* stems identified as the main components the hexadecanoic acid ethyl ester, the octadecanoic acid ethyl ester and phytol ^30^.

Fourteen alkanes were identified with a hydrocarbon chain ranging from C20 to C35. All of them has been previously identified as a constituents of plant stems by GC-MS analysis ^31^ but just some of them has been identified in plants of the genus *Cissus*. In *C. quadrangularis* stems was reported the presence of pentacosane, heptacosane, pentatriacontane, and hentriacontane ^32^. In *C. vitigenea* was reported docosane, heptacosane and nonadecane ^32^, whereas the nonacosane in *C. cornifolia* ^33^, the cyclooctacosane was described by GC-MS in *C. sicyoides* ^34^. In accordance, the heptacosane, nonacosane and hentriacontane, are predominant cuticular wax components in plants ^16^.

The carboxylic acids founded were the n-hexadecanoic which is also the principal component of the hexane extract of the stems of *C. quadrangularis* ^30^ and their aerial parts (aqueous alcoholic extract) ^35^. Palmitic acid is also present in *C. vitiginea* and their content in plant extracts is highest in low polarity solvents ^32^. Their role is as structural lipid, with influence in fluidity of cellular membranes of higher plants ^36^. In the case of the n-hexadecanoic acid ethyl ester, it was found as the major constituent of hexane extract from roots in *C. quadrangularis* ^30^, and it’s a common fatty acid with methyl branch in plant waxes ^20^. Other two saturated fatty acids; the octadecanoic acid, also called stearic acid, is a major constituent in the hexane extract from the stems of *C. quadrangularis*, and in the methanolic extract of the entire plant ^30, 37^. This fatty acid is one of the most abundant plant membrane fatty acids and play structural roles ^38^. In the case of eicosanoic acid, is also a major constituent of hexane extract of roots in *C. quadrangularis* ^30^, but in general is a minor constituent of plant cell membranes ^20^. The unsaturated fatty acids present in the hexane extract of *C. trifoliata* were the (Z,Z) 9,12-octadiecadienoic acid and (Z)-9-octadecenoic acid. The former was reported for the first time in the methanolic extract of *C. quadrangularis* stems ^39^ and the last one as a major constituent of hexane extracts roots in the same species ^30^. Both fatty acids, the linoleic acid and the oleic acid play roles in increasing the fluidity of plant membranes ^38^. This is the first report of the presence of the alcohol 1,30-triacontanediol in *Cissus* species, however is a common solid alcohol from plants present in their cuticular wax ^16^ acting as a growth regulator ^40^.

Among the terpenes identified, the squalene is the simplest and the most abundant, according with previous studies of the hexane extract of the stems from *Cissus* plants ^13^. On the other hand, the lupeol was the only pentacyclic terpene identified by GC-MS, but is also present as a major constituent of hexane extract of stems and roots in *C. quadrangularis* ^13, 30^. Its biosynthesis is induced by pathogens, and exerts antimicrobial activities ^41^. In the case of the triterpenes, seven sterols were identified; β-sitosterol, stigmasta-5,22-dien-3-ol, stigmast-4-en-3-one, campesterol, brassicasterol and epicholesterol. The first two were obtained as mixture in compound 4 as previously discussed, in the case of campesterol it was isolated from hexane ^30^ and ethanolic extract of *C. quadrangularis* stems ^39^ and is also found in methanolic extract of the roots of *C. rheifolia* ^26^. Stigmast-4-en-3-one was previously isolated from the ethanolic extract of *C. quadrangularis* stems ^42^. Although all sterols share a structural role in plant membranes fluidity ^27^, changes in plant sterol structure, such as the observed in stigmast-4-en-3-one is associated with infestation by insects, acting as antifeedant ^43^. Brassicasterol and epicholesterol has been reported by GC-MS analysis of *Cissus* plants but in minor concentrations, since are abundant in algae and mammals respectively ^30^.

Finally, the GC-MS analysis also shows the presence of bis (2-ethylhexyl) phthalate, and di (2-ethylhexyl) isophthalates, as a major constituent (∼3%) of the stems of *C. trifoliata*, however they are not the product of biosynthesis in plants. To date, no genes are related with the production of this type of molecules in plants, but recently, they were characterized in filamentous fungi, thus they may be related with the metabolism of endophytes in *Cissus* plants ^44^. On the other hand, those compounds are generally reported as a common contaminant in almost all laboratory equipment and reagents such as plastics, glassware, aluminum foil, cork, rubber, glass wool, and solvents ^45^. However, studies of chemistry by GC-MS from *Cissus quadrangularis, vitigenea*, and *aralioides* also report phtlates as major constituents in the extracts. Thus seem plausible that compounds reported as phthalates by GC-MS techniques are false spectral match identifications due structural relationship with phytocompounds ^46^.

**Figure 3.**
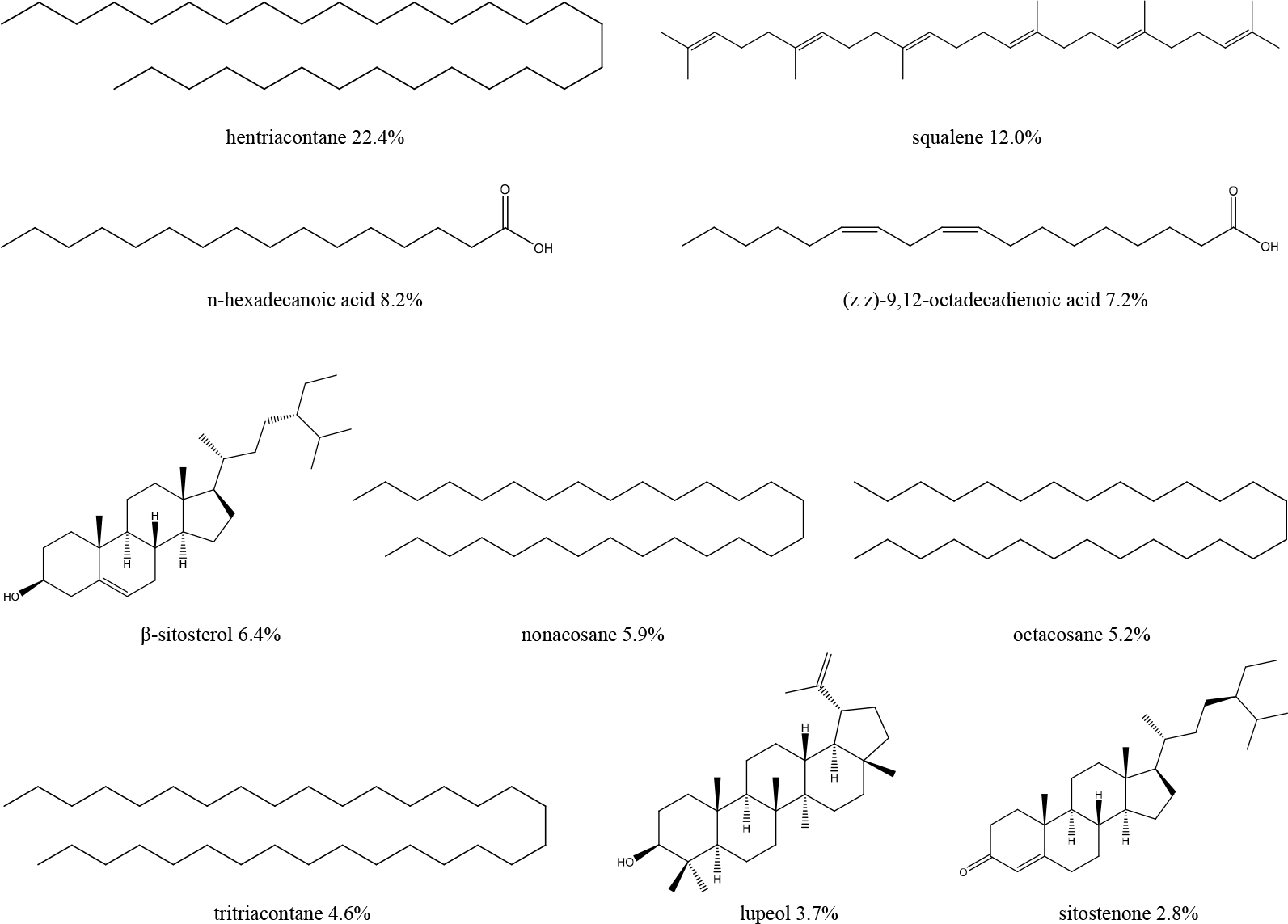
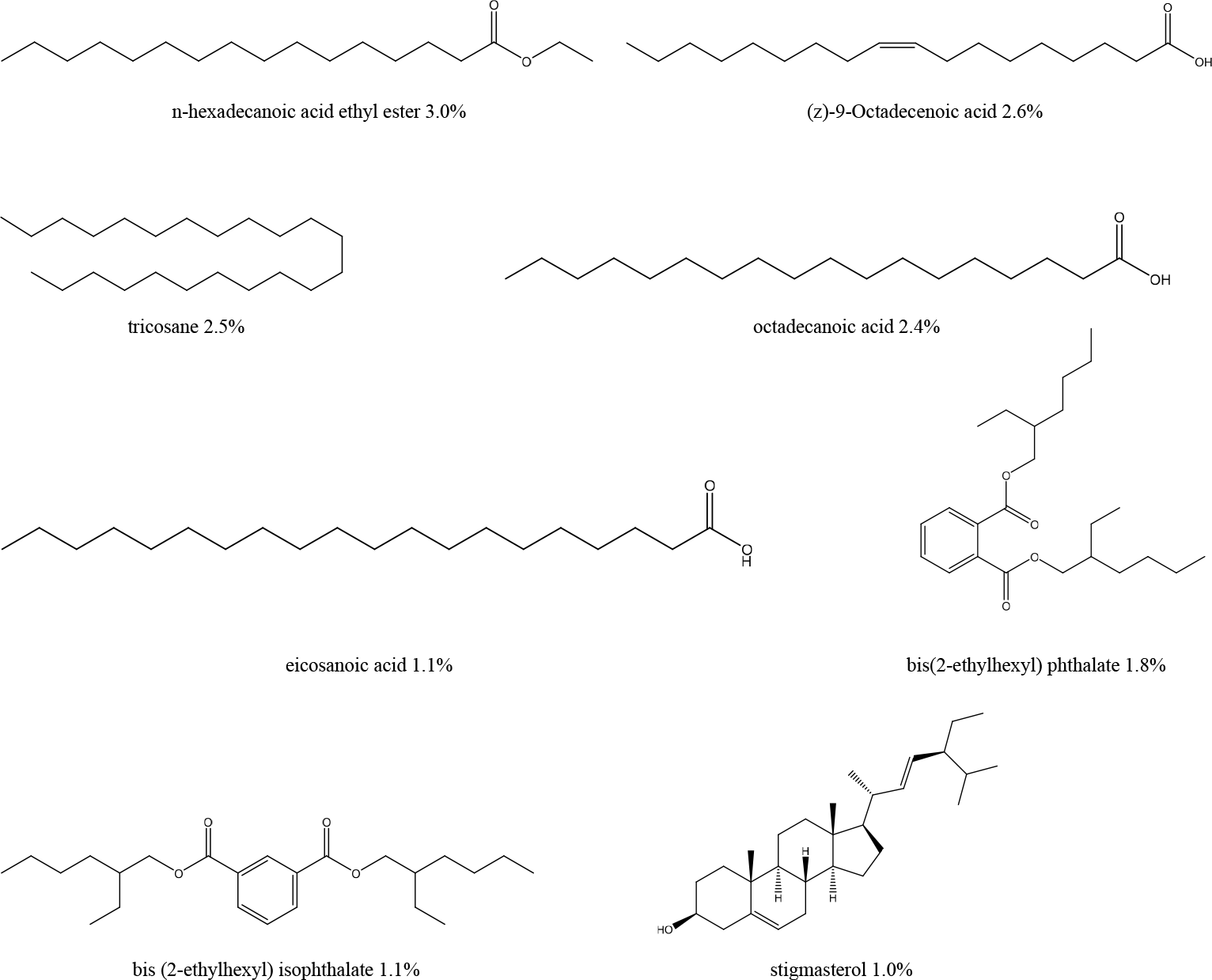
Chemical structures of most abundant compounds in the hexane extract of *C. trifoliata* stems according to GC-MS analysis

## Conclusion

The hexane extract of *C. trifoliata* stems presents five major compounds which spectroscopic signals corresponds with higher plant n-alkanes such as hentriacontane and octatetracontane, with a long-chain fatty acid, the octacosanoic acid, a wax ester composed of an alcohol and a fatty acid, the tricosyl tetracosanoate and a mixture of the main plant sterols, the β-sitosterol and stigmasterol. The GC-MS analysis identified 35 compounds and the most abundant were hentriacontane, squalene, palmitic acid, oleic acid, β-sitosterol, and nonacosane. Most of them were previously reported in *Cissus* plants except for heneicosane, tricosane, hexacosene, and tritiacontane. Overall, the phytochemical content of the hexane extract of *C. trifoliata* stems corresponds mainly to higher alkanes, fatty acids and sterols, which are the major components of cuticular waxes and plant cell membranes.

## Author Contributions

L.F.M.L prepared the extract, established the chromatographic conditions, participated in the GC-MS and NMR analysis, and wrote the first manuscript draft. B.A.E.R and J.L.G.L analyzed the spectrometric and spectroscopic data. I.B.R and M.R.C.C contributed to the design and supervised the development of this project. All authors contributed to a critical reading of the manuscript.

## Acknowledgments

The manuscript was taken in part from the PhD thesis of LFML. The principal author thanks CONACYT-Mexico for the scholarship (210600) to carry out his PhD studies.

## References

1. Standley, P. C. Trees and shrubs of Mexico. Smithsonian Institution: 1967; Vol. 1.

2. McCartney, P. SEINet: metadata-mediated access to distributed ecological data. LTER. DataBits Spring 2003, 1.

3. Heinrich, M.; Ankli, A.; Frei, B.; Weimann, C.; Sticher, O. Medicinal plants in Mexico: Healers’ consensus and cultural importance. Social Science & Medicine 1998, 47 (11), 1859–1871.

4. de las Mercedes Rodríguez, L. Etnobotánica maya: Algunas plantas de uso medicinal en estomatología. Revista ADM 2015, 72 (1).

5. Estrada-Castillón, E.; Soto-Mata, B. E.; Garza-López, M.; Villarreal-Quintanilla, J. Á.; Jiménez-Pérez, J.; Pando-Moreno, M.; Sánchez-Salas, J.; Scott-Morales, L.; Cotera-Correa, M., Medicinal plants in the southern region of the State of Nuevo León, México. Journal of ethnobiology and ethnomedicine 2012, 8 (1), 45.

6. Méndez-López, L. F.; Garza-González, E.; Ríos, M. Y.; Ramírez-Cisneros, M. Á.; Alvarez, L.; González-Maya, L.; Sánchez-Carranza, J. N.; Camacho-Corona, M. d. R. Metabolic profile and evaluation of biological activities of extracts from the stems of Cissus trifoliata. International Journal of Molecular Sciences 2020, 21 (3), 930.

7. Méndez-López, L. F.; Caboni, P.; Arredondo-Espinoza, E.; Carrizales-Castillo, J. J.; Balderas-Rentería, I.; Camacho-Corona, M. d. R. Bioassay-guided identification of the antiproliferative compounds of Cissus trifoliata and the transcriptomic effect of resveratrol in prostate cancer PC3 cells. Molecules 2021, 26 (8), 2200.

8. Ahmadu, A.; Onanuga, A.; Aquino, R. Flavonoid glycosides from the leaves of Cissus ibuensis hook (vitaceae). African Journal of Traditional, Complementary and Alternative Medicines 2010, 7 (3).

9. Jain, V.; Thakur, A.; Hingorani, L.; Laddha, K. Lipid constituents from Cissus quadrangularis leaves. Pharmacognosy Research 2009, 1 (4), 231.

10. Laitonjam, W. S.; Yumnam, R. S.; Kongbrailatpam, B. D. Study on isolation and comparison of the chemical compositions of Cissus adnata Roxb. leaves and Smilax lanceaefolia Roxb. roots and their free radical scavenging activities. International Research Journal of Pure and Applied Chemistry 2011, 1 (1), 1.

11. Adesanya, S. A.; Nia, R.; Martin, M. T.; Boukamcha, N.; Montagnac, A.; Païs, M. Stilbene derivatives from Cissus quadrangularis. Journal of Natural Products 1999, 62 (12), 1694–1695.

12. Thakur, A.; Jain, V.; Hingorani, L.; Laddha, K. Phytochemical studies on Cissus quadrangularis Linn. J Pharmacognosy Research 2009, 1 (4), 213.

13. Pathomwichaiwat, T.; Ochareon, P.; Soonthornchareonnon, N.; Ali, Z.; Khan, I. A., Prathanturarug, S. Alkaline phosphatase activity-guided isolation of active compounds and new dammarane-type triterpenes from Cissus quadrangularis hexane extract. Journal of ethnopharmacology 2015, 160, 52–60.

14. Gupta, M. M.; Verma, R. K. Lipid constituents of Cissus quadrangularis. Phytochemistry 1991, 30 (3), 875–878.

15. Cookson, D. J.; Smith, B. E. Determination of structural characteristics of saturates from diesel and kerosine fuels by carbon-13 nuclear magnetic resonance spectrometry. Analytical Chemistry 1985, 57 (4), 864–871.

16. Lara, I.; Belge, B.; Goulao, L. F. A focus on the biosynthesis and composition of cuticle in fruits. Journal of agricultural and food chemistry 2015, 63 (16), 4005–4019.

17. Singh, N. K.; Singh, V. Phytochemistry and pharmacology of Ichnocarpus frutescens. Chinese journal of natural medicines 2012, 10 (4), 241–246.

18. Lim, T. K., Edible medicinal and non-medicinal plants. Springer: 2012; Vol. 1.

19. Aggarwal, B.; Ali, M.; Singh, V.; Singla, R. K. Isolation and characterization of phytoconstituents from the stems of Ichnocarpus frutescens. Chinese Journal of Natural Medicines 2010, 8 (6), 401–404.

20. Gunstone, F. D.; Harwood, J. L.; Dijkstra, A. J. The lipid handbook with CD-ROM. CRC press: 2007.

21. Harwood, J., Lipid metabolism. In The lipid handbook, Springer: 1986; pp 485–525.

22. Chaturvedula, V. S. P.; Prakash, I. Isolation of Stigmasterol and β-Sitosterol from the dichloromethane extract of Rubus suavissimus. 2012.

23. Xie, Y.; Deng, P.; Zhang, Y.; Yu, W. Studies on the chemical constituents from Cissus assamica. Zhong yao cai= Zhongyaocai= Journal of Chinese medicinal materials 2009, 32 (2), 210–213.

24. Sani, Y.; Musa, A.; Tajuddeen, N.; Abdullahi, S.; Abdullahi, M.; Pateh, U.; Idris, A. Isoliquiritigenin and-sitosterol from Cissus polyantha Tuber Glig and Brandt. Journal of Medicinal Plants Research 2015, 9 (35), 918–921.

25. Pan, G.; Li, W.; Luo, P.; Qin, J.; Su, G. Study on steroidal and triterpenoid constituents from Cissus pteroclada. Zhong yao cai= Zhongyaocai= Journal of Chinese medicinal materials 2013, 36 (8), 1274–1277.

26. Saifah, E.; Vaisiriroj, V.; Kelley, C. J.; Higuchi, Y. Constituents of the roots of Cissus rheifolia. Journal of Natural Products 1987, 50 (2), 328–328.

27. Griebel, T.; Zeier, J. A role for β-sitosterol to stigmasterol conversion in plant–pathogen interactions. The plant journal 2010, 63 (2), 254–268.

28. Toro-Vazquez, J. F.; Mauricio-Pérez, R.; González-Chávez, M. M.; Sánchez-Becerril, M.; de Jesús Ornelas-Paz, J.; Pérez-Martínez, J. D. Physical properties of organogels and water in oil emulsions structured by mixtures of candelilla wax and monoglycerides. Food research international 2013, 54 (2), 1360–1368.

29. Heredia-Guerrero, J. A.; Benítez, J. J.; Domínguez, E.; Bayer, I. S.; Cingolani, R.; Athanassiou, A.; Heredia, A. Infrared and Raman spectroscopic features of plant cuticles: a review. Frontiers in plant science 2014, 5, 305.

30. Vishnuthari., N.; Sripathi., S. K. GC-MS analysis of hexane extract of stems and roots of the ethnomedicinal plant Cissus quadrangularis Linn. J. Chem. Bio. Phy. Sci. 2015, 5 (4), 3954–3963.

31. Silva, G. C.; Bottoli, C. B. Analyses of Passiflora compounds by chromatographic and electrophoretic techniques. J Critical Reviews in Analytical Chemistry 2015, 45 (1), 76–95.

32. Rosy, B. A.; Rosakutty, P. GC-MS analysis of methanol wild plant and callus extracts from three Cissus species, Family Vitaceae. Journal of chemical and pharmaceutical research 2012, 4 (7), 3420–3426.

33. Chipiti, T.; Ibrahim, M. A.; Koorbanally, N. A.; Islam, M. S. In vitro antioxidant activity and GC-MS analysis of the ethanol and aqueous extracts of Cissus cornifolia (Baker) Splanch (Vitaceae) parts. ActaPoloniaePharmaceutica Drug Research 2015, 72 (1), 119–127.

34. Beltrame, F. L.; Sartoretto, J. L.; Bazotte, R. B.; Cuman, R. N.; Cortez, D. A. G.; Fernandes, L. C.; Tchaikovski, O. Phytochemical study and evaluation of the antidiabetic potential of Cissus sicyoides L.(Vitaceae). Química Nova 2001, 24 (6), 783–785.

35. Kumar, S.; Anandan, A.; Jegadeesan, M. Identification of chemical compounds in Cissus quadrangularis L. Variant I of different samples using GC-MS analysis. Archives of Applied Science Research 2012, 4 (4), 1782–1787.

36. Zhukov, A. Palmitic acid and its role in the structure and functions of plant cell membranes. Russian journal of plant physiology 2015, 62 (5), 706–713.

37. Eswaran, R.; Anandan, A.; Doss, A.; Sangeetha, G.; Anand, S. Analysis of chemical composition of Cissus quadrangularis linn by GC-MS. Asian j pharm clin res 2012, 2, 139–40.

38. Heldt, H. Plant biochemistry. Elsevier Academic Press. XXIV: 2005.

39. Chanda, S.; Baravalia, Y.; Nagani, K. Spectral analysis of methanol extract of Cissus quadrangularis L. stem and its fractions. Journal of Pharmacognosy and Phytochemistry 2013, 2 (4).

40. Koonce, S. D.; Brown, J. A study of the alcohols of carnauba wax. Journal of the American Oil Chemists’ Society 1944, 21 (8), 231–234.

41. Wal, P.; Wal, A.; Sharma, G.; Rai, A. Biological activities of lupeol. Systematic Reviews in Pharmacy 2011, 2 (2), 96.

42. Sheikh, S.; Siddiqui, S.; Dhasmana, A. Cissus quadrangularis Linn. Stem ethanolic extract liberates reactive oxygen species and induces mitochondria mediated apoptosis in KB cells. Pharmacognosy magazine 2015, 11 (Suppl 3), S365.

43. Jing, X.; Grebenok, R. J.; Behmer, S. T. Plant sterols and host plant suitability for generalist and specialist caterpillars. Journal of insect physiology 2012, 58 (2), 235–244.

44. Tian, C.; Ni, J.; Chang, F.; Liu, S.; Xu, N.; Sun, W.; Xie, Y.; Guo, Y.; Ma, Y.; Yang, Z. J. S. Bio-Source of di-n-butyl phthalate production by filamentous fungi. 2016, 6, 19791.

45. Humans, I. W. Some chemicals present in industrial and consumer products, food and drinking-water. IARC monographs on the evaluation of carcinogenic risks to humans 2013, 101, 9.

46. Koo, I.; Kim, S.; Zhang, X. J. Comparative analysis of mass spectral matching-based compound identification in gas chromatography–mass spectrometry. 2013, 1298, 132–138.

